# Vascular waveform analysis using Bayesian pulse deconvolution

**DOI:** 10.64898/2026.02.09.699383

**Authors:** Parker S. Ruth, Tommy DeBenedetti, Lily O’Brien, James A. Landay, Todd P. Coleman, Emily B. Fox

## Abstract

Vascular waveforms, which measure bulk flow in blood vessels, are widely used to measure vital signs, diagnose conditions, and predict long-term health outcomes. Analyzing vascular waveforms depends on three fundamentally interdependent tasks: signal filtering, pulse timing detection, and pulse shape extraction. We hypothesized that Bayesian pulse deconvolution can achieve improved performance on all three tasks by solving them jointly. This method uses an analytical, generative model of vascular waveforms with priors informed by physical and biological domain knowledge. In simulations, Bayesian pulse deconvolution achieves better performance on all tasks compared with existing algorithms: 90% reduction of median filtering error, 60% reduction in pulse timing error, and 85% reduction in shape extraction error. The advantages in simulations extend to human recordings of photoplethysmography waveforms. Taking real time-synchronized electrocardiogram R-R intervals as a proxy ground truth, Bayesian pulse deconvolution achieves 40% lower pulse interval estimation error (RMSE = 5.1 ms) compared with typical algorithms (RMSE = 8.3 ms, p=1e–10). By extracting more accurate and informative insights from vascular waveforms, Bayesian pulse deconvolution could advance a wide array of health technologies that rely on interpreting signals from blood vessels.

## INTRODUCTION

Vascular waveforms, which measure bulk fluid transport in blood vessels, are among the most ubiquitous and important signals recorded from the human body. These signals, including intravascular blood pressure, blood flow velocity, photoplethysmography, and applanation tonometry, are readily acquired using invasive hemodynamic monitors, ^1–3^ non-invasive wearable sensors, ^4–6^ and a host of emerging sensors. ^7^ Vascular waveforms, alone and in combination with other measures, are widely used to monitor health and detect disease. They give rise to many important vital signs, including heart rate variability, ^8^ blood oxygenation, ^9^ blood pressure, ^10–12^ blood volume, ^13^ and cardiorespiratory fitness. ^14^ Vascular waveforms can reveal cardiovascular health problems such as arrhythmia, ^15^ hemodynamic decompensation, ^16^ and atherosclerosis, ^17^ and they can even aid in predicting significant health outcomes such as heart attack, stroke, and all-cause mortality. ^18,19^

All of these applications require clinically interpreting the rhythms and shapes of heart beats in sensor data. This involves three fundamental signal processing tasks: filtering noisy signals, detecting pulse timing, and extracting pulse shapes. In current practice, these tasks are typically performed in a sequential pipeline. ^20–23^

1. **Filtering noisy signals** is important for isolating vascular waveforms from sensor error, baseline drift, artifacts, and other non-physiological sources. This may be performed by linear digital filters, ^24–29^ Savitzky-Golay filters, ^29^ adaptive comb filters, ^30,31^ or neural networks. ^32^
2. **Detecting pulse timing** is important for extracting beat-to-beat intervals, heart rate, or pulse transit times. Pulse timing is typically measured by local extrema of the 0th, 1st, and 2nd derivatives of filtered vascular signals. ^33–35^
3. **Extracting pulse shapes** is important for calculating health metrics such as blood oxygenation, arterial stiffness, and blood pressure. This is typically performed by segmentation and pulse averaging. ^22,23^

The choice of algorithm for each step has consequences that propagate through the waveform analysis pipeline, impacting the accuracy of downstream applications such as measuring blood pressure ^26^ or blood oxygenation. ^30^ For example, Chebyshev type-II filters score highly on a signal quality heuristic ^27^ but actually introduce phase distortions that significantly impact extracted pulse shapes. ^36^ Similarly, the choice of heuristic timing markers impacts the estimated heart beat timing and segmentation. ^37^ In fact, filtering, timing detection, and shape extraction are fundamentally interdependent problems. Although current algorithms treat them independently, we hypothesize that optimally solving any one problem requires solving all three problems jointly.

Designing supervised algorithms for vascular waveforms analysis is complicated by a lack of ground truth data for training and testing. Every sensor measurement has some noise, so there is no *bona fide* ground truth for de-noising filters. Instead, filters are currently optimized based on heuristic measures of signal quality, which may distort true physiology. ^38,39^ While electrocardiography (ECG) is the gold standard for measuring pulse timing, the time delay between ECG pulses and vascular pulse onset is variable between individuals and between heart beats, ^40^ making ECG an inexact source of pulse timing labels. Furthermore, there is no ground truth method for isolating the individual shapes of potentially overlapping pulses in vascular waveforms. Inadequate ground truth presents a fundamental obstacle for data-driven algorithms that rely on trusted labels for training and testing. Instead, to overcome inadequate training data, we propose encoding physical and biological domain knowledge in an unsupervised algorithm. To overcome inadequate testing labels, we evaluate algorithms with a combination of simulated data and human recordings of vascular waveforms.

To jointly solve the filtering, timing, and shape problems, we propose a unified Bayesian pulse deconvolution algorithm that encodes domain knowledge in its constraints and priors, and quantifies uncertainty in its solutions. We model pulse timing as a semi-periodic point process, informed by the biology of heart rhythms. We model a vascular waveform as a convolution of the pulse times with an impulse response, informed by the physical principle of wave superposition. ^41^ We combine data and domain knowledge to design Bayesian prior densities over unobservable variables in the model and compute Bayesian posterior densities conditioned on observed signals.

This work presents a unified approach to vascular waveform analysis. Previously, deconvolution methods have been proposed for analyzing other sparse or semi-periodic biosignals such as neural activity ^42^ and blood oxygenation level dependent functional magnetic resonance. ^43^ Some related work has proposed generative AI models for cardiovascular biosignals. ^44^ Other related work has proposed non-Bayesian iterative algorithms for adaptive template-matching filters ^45^ or for decomposing combined cardiac and vascular waveforms into impulse trains and impulse responses. ^46,47^ However, these methods lack domain-grounded priors for vascular waveforms. Probabilistic models have been proposed for simulating synthetic vascular waveforms ^48–50^; however, these models are not suitable for processing human recordings with Bayesian inference. We instead propose a Bayesian pulse deconvolution algorithm to jointly solve filtering, timing detection, and shape extraction on vascular waveforms. We use simulated data to systematically evaluate the advantages of Bayesian pulse deconvolution over representative existing algorithms. Finally, we test whether the simulated results extend to human recordings of pulse timing using raw vascular waveforms with time-synchronized electrocardiogram signals.

## RESULTS

We compared the accuracy of Bayesian pulse deconvolution and traditional algorithms for signal filtering, pulse timing detection, and pulse shape extraction. In the absence of acceptable ground truth labels for these tasks, we first evaluate the algorithms on 100 simulated eleven-second waveforms sampled at 64 Hz with –20dB Gaussian noise. These simulations let us evaluate against the original noiseless signals, exact pulse times, and isolated pulse shapes. We then showcase the application of Bayesian pulse deconvolution to an experimental dataset of human recordings.

### Signal filtering

Vascular waveforms are typically filtered with linear time-invariant filters, which are characterized by their frequency-dependent transfer functions. These filters assume that signal and noise are separable in the frequency domain, which is generally untrue (for example, white Gaussian noise has uniform power across all frequencies). Thus, linear filters either incompletely attenuate noise or partially attenuate desired signals. Furthermore, since commonly used infinite impulse response filters introduce phase delays in their pass bands, they can cause signal distortions that supersede their noise attenuation (**Fig. 1a**). Instead, Bayesian pulse deconvolution recovers noiseless signals by finding maximum *a posteriori* estimates of an analytical model (**Fig. 1b**), making no assumptions about frequency-domain noise separation.

**Fig. 1.**
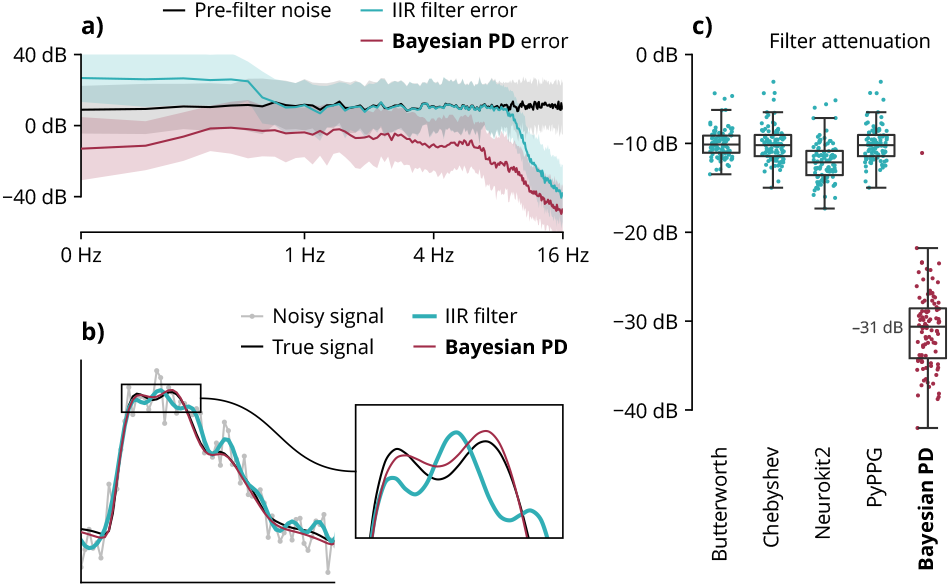
Signal filtering accuracy in simulation. (**a**) A typical 2nd-order 0.5-12Hz Butterworth bandpass filter incompletely attenuates noise and introduces phase distortions in the physiological frequency range. Bayesian pulse deconvolution (PD) achieves greater noise attenuation across all frequencies. Shaded regions represent ±1 standard deviation across 100 simulations. (**b**) Bayesian pulse deconvolution (PD) recovers the waveform with less filter distortion than a typical filter. (**c**) Across 100 simulated waveforms, Bayesian pulse deconvolution achieves the best noise attenuation. Each point represents one simulated waveform. (Baseline algorithms are defined in Table 1.)

We compared the noise attenuation of Bayesian pulse deconvolution against baselines representing “optimal” filters reported in prior literature ^27,36^ and standard published vascular waveform analysis algorithms. ^21,22^ Across 100 simulations, the traditional filters achieved median noise attenuation of 10 to 12 dB, while the Bayesian algorithm achieved a median noise attenuation of 31 dB, corresponding to a 90% reduction in the error signal power (**Fig. 1c**). The filtering performance improvement holds across signal durations, sampling rates, and noise levels **(Sup. Fig. 1)**.

**Table 1.**
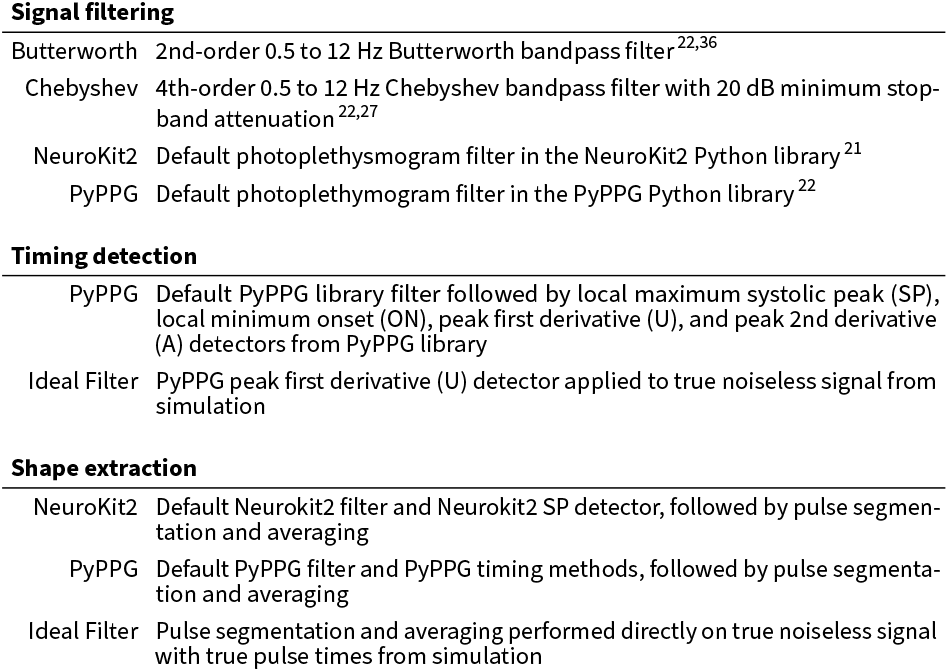
Baseline algorithms. Descriptions of baseline vascular waveform analysis algorithms compared against Bayesian pulse deconvolution.

Signal filtering
|  |  |
| --- | --- |
| Butterworth | 2nd-order 0.5 to 12 Hz Butterworth bandpass filter<sup>22,36</sup> |
| Chebyshev | 4th-order 0.5 to 12 Hz Chebyshev bandpass filter with 20 dB minimum stop-band attenuation<sup>22,27</sup> |
| NeuroKit2 | Default photoplethysmogram filter in the NeuroKit2 Python library<sup>21</sup> |
| PyPPG | Default photoplethysmogram filter in the PyPPG Python library<sup>22</sup> |
| Timing detection |  |
| PyPPG | Default PyPPG library filter followed by local maximum systolic peak (SP), local minimum onset (ON), peak first derivative (U), and peak 2nd derivative (A) detectors from PyPPG library |
| Ideal Filter | PyPPG peak first derivative (U) detector applied to true noiseless signal from simulation |

|  |  |
| --- | --- |
| NeuroKit2 | Default Neurokit2 filter and Neurokit2 SP detector, followed by pulse segmentation and averaging |
| PyPPG | Default PyPPG filter and PyPPG timing methods, followed by pulse segmentation and averaging |
| Ideal Filter | Pulse segmentation and averaging performed directly on true noiseless signal with true pulse times from simulation |

### Pulse timing detection

Typically, pulse times are detected by heuristic markers on filtered signals. Common heuristic markers are local extrema of the 0th, 1st, or 2nd derivatives of the signal (**Fig. 2a**). These markers are sensitive to measurement noise and filter distortions because they only use local features of the pulse shape. By contrast, Bayesian pulse deconvolution uses information from the entire waveform to determine pulse timing, while accounting for overlap between adjacent pulses (**Fig. 2b**).

**Fig. 2.**
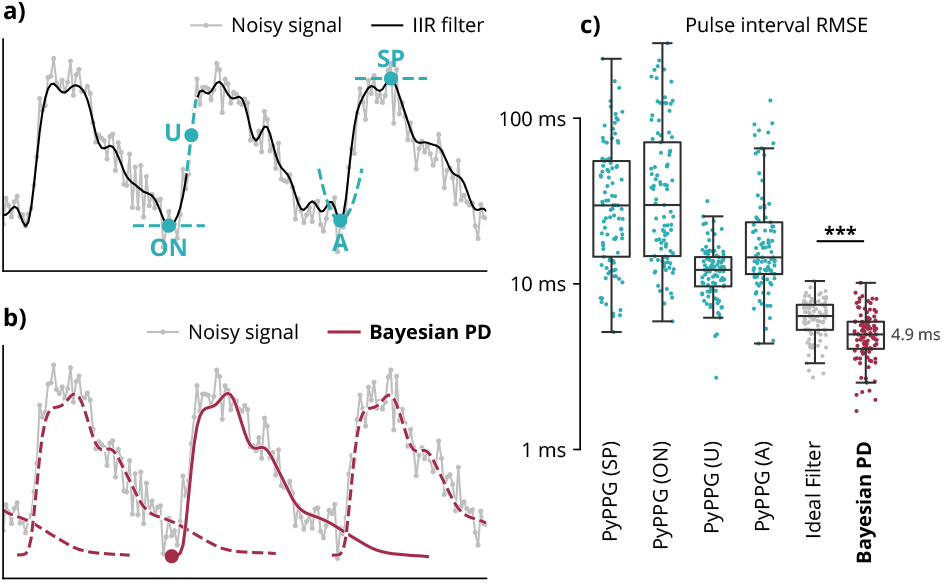
Timing detection accuracy in simulation. (**a**) Typical pulse timing features are extracted from heuristic positions on a pre-filtered signal, such as the local maximum systolic peak (SP), local minimum onset (ON), peak first derivative (U), or peak 2nd derivative (A). (**b**) Bayesian pulse deconvolution (PD) aligns timing to the entire heartbeat cycle. (**c**) Across 100 simulated waveforms, Bayesian pulse deconvolution achieves the best root-mean-square error (RMSE). The “Ideal Filter” baseline takes the peak first derivative (U) from the true noiseless waveforms from simulation. This baseline has a higher median RMSE than Bayesian PD (p=1e–5). Each point represents one simulated waveform. (Baseline algorithms are defined in Table 1.)

In 100 simulations, we estimated pulse times and measured resulting beat-to-beat intervals using Bayesian pulse deconvolution as well as four common heuristic markers as implemented by the PyPPG Python package. ^22^

- SP: local maximum of signal (systolic peak)
- ON: local minimum of signal (onset)
- U: local maximum of first derivative (velocity)
- A: local maximum of second derivative (acceleration)

As reported in prior literature, ^33,37^ the peak first derivative (U) most accurately measures beat-to-beat intervals (median RMSE = 12.0 ms). Bayesian pulse deconvolution reduces the error by 60% (median RMSE = 4.9 ms) (**Fig. 2c**). The pulse timing detection improvement holds across signal durations, sampling rates, and noise levels **(Sup. Fig. 1)**.

The accuracy of heuristic marker timing may be impacted by upstream filter distortion. To differentiate error due to the heuristic markers from error due to filter distortion, we extracted the peak first derivative from the ideal noiseless signal. In this case, the heuristic marker still has higher error (RMSE = 6.4 ms) than Bayesian pulse deconvolution (p=1e–5).

### Pulse shape extraction

Typically pulse shape is extracted by segmenting waveforms based on heuristic pulse timing markers (**Fig. 3a**). By averaging pulses, this approach can achieve substantial noise attenuation. However, error in heuristic pulse times can impact extracted shape resolution. Furthermore, overlap between adjacent heartbeats causes distortion and truncation of the extracted pulse shape. By contrast, Bayesian pulse deconvolution recovers the full underlying pulse shape as an analytical function, while accounting for overlapping pulses (**Fig. 3b**).

**Fig. 3.**
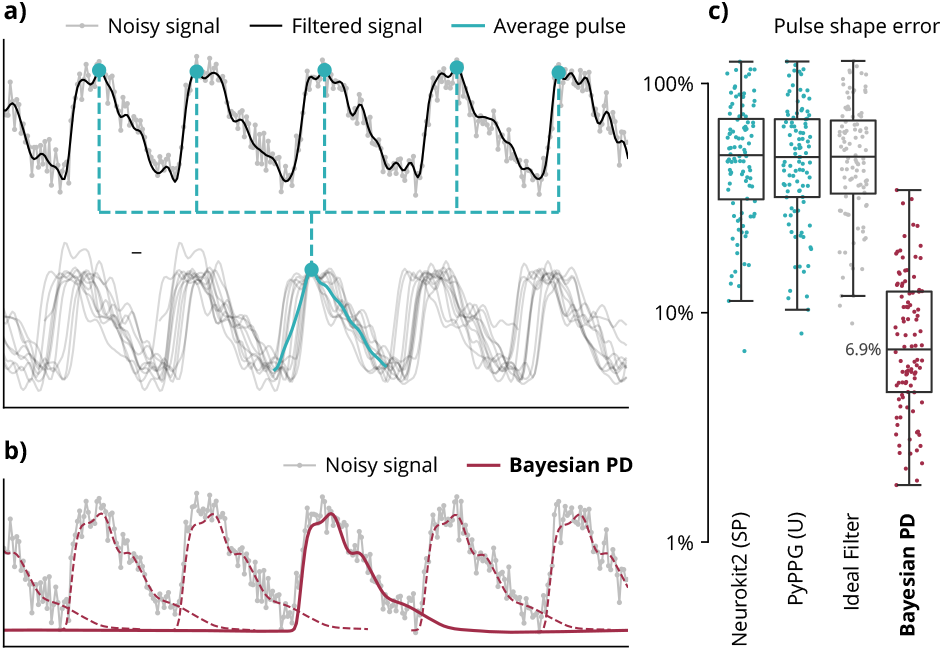
Shape extraction accuracy in simulation. (**a**) Typically, pulse shapes are extracted by aligning and averaging a filtered signal based on heuristic timing markers, ignoring overlaps between pulses. (**b**) Bayesian pulse deconvolution (PD) explicitly models overlaps in pulse shapes. (**c**) Across 100 simulated waveforms, Bayesian pulse deconvolution achieves the lowest percent error of the pulse shape area. The “Ideal Filter” baseline is the average pulse shape from the true noiseless signal using the true exact pulse times from simulation. Each point represents one simulated waveform. (Baseline algorithms are defined in Table 1.)

We assessed the accuracy of pulse shape extraction based on the total percent area of error. This metric (defined in Methods below) captures error in the contours of the pulse as well as erroneous truncation of the pulse. Across 100 simulations, typical pulse shape extraction methods achieved a median error of 48-49%, while Bayesian pulse deconvolution achieved a median error of 6.9%, corresponding to a 85% reduction in shape extraction error (**Fig. 3c**). The pulse shape extraction improvement holds across signal durations, sampling rates, and noise levels **(Sup. Fig. 1)**.

The accuracy of the traditional pulse shape extraction method may be impacted by upstream filtering and pulse timing error. To isolate the error solely due to the segmentation and averaging, we applied shape extraction to an ideal noiseless signal and exact pulse times from the simulation. In this case, median extracted shape error was not improved over the other baselines, indicating that error is primarily due to the pulse overlaps and truncations inherent in this shape extraction method.

### Evaluations on human recordings

We sought to use human recordings to test whether Bayesian pulse deconvolution’s advantages in simulations translate into better performance on real data. We evaluated the ability to estimate gold standard, ECG-based R-R intervals from wrist-worn photoplethysmogram waveforms (**Fig. 4a**), a task essential for wearable heart rate monitoring and arrhythmia detection. ^51,52^ Bayesian pulse deconvolution can detect sub-sampling-rate changes in heart beat timing such as low-amplitude sinus arrhythmia patterns that traditional methods cannot accurately capture (**Fig. 4b**). We compared the best-performing traditional pulse timing method and Bayesian pulse deconvolution on 105 recordings from 21 individuals in supine, seated, standing, walking, and running conditions (**Fig. 4c**). While the traditional pulse timing method achieved a median RMSE of 8.3 ms, Bayesian pulse deconvolution achieved a median RMSE of 5.1 ms, constituting a 40% error reduction.

**Fig. 4.**
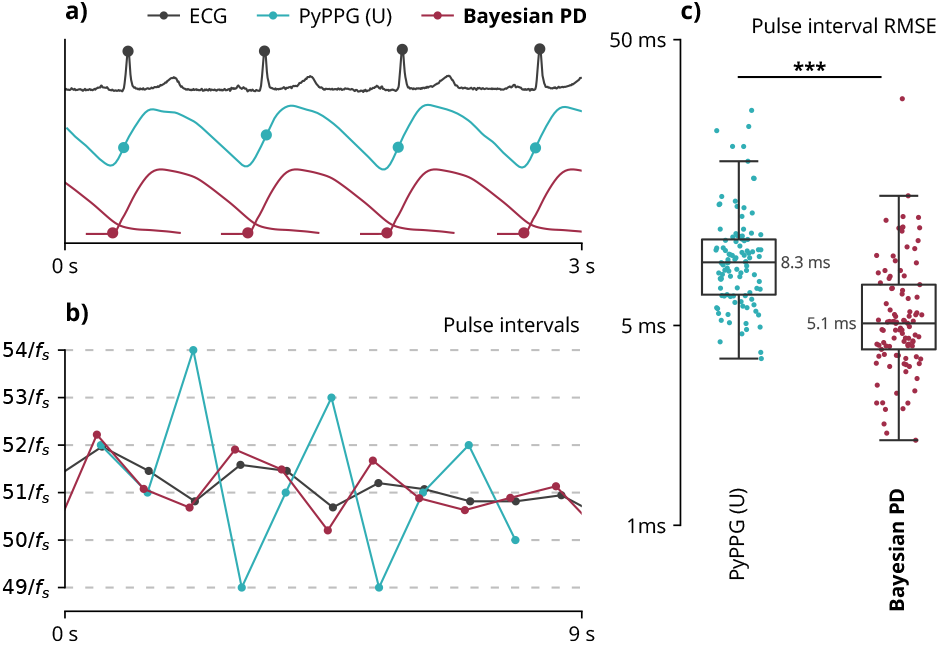
Pulse timing evaluation on human recordings. (**a**) Pulse timing analysis of experimental recordings from a 48-year-old supine participant. Peaks of the electrocardiogram (ECG) are a proxy ground truth for evaluating the accuracy of pulse intervals extracted from a typical PyPPG (U) algorithm (defined in Table 1) and Bayesian pulse deconvolution (PD). (**b**) Pulse intervals extracted from the waveforms above. Dashed gray lines represent integer multiples of the sample period 1/*f*_*s*_. (**c**) Across 105 human recordings, Bayesian pulse deconvolution has a lower median root-mean-square error (RMSE) relative to ECG (p=1e–10). Each point represents one vascular waveform. Values are on a logarithmic scale.

### Uncertainty quantification

In addition to producing maximum *a posteriori* solutions, a Bayesian framework also allows us to compute the full joint posterior distribution that describes uncertainty over the solution space. Posterior uncertainty responds as expected to sampling rate and signal-to-noise ratio (SNR). A signal with high sampling rate and high SNR has high certainty (**Fig. 5a**). A signal with either low sampling rate (**Fig. 5b**) or low SNR (**Fig. 5c**) has moderate certainty. A signal with both low sampling rate and low SNR has low certainty (**Fig. 5d**). Systematic evaluation of algorithm performance across sampling parameters is shown in **Sup. Fig. 1**.

**Fig. 5.**
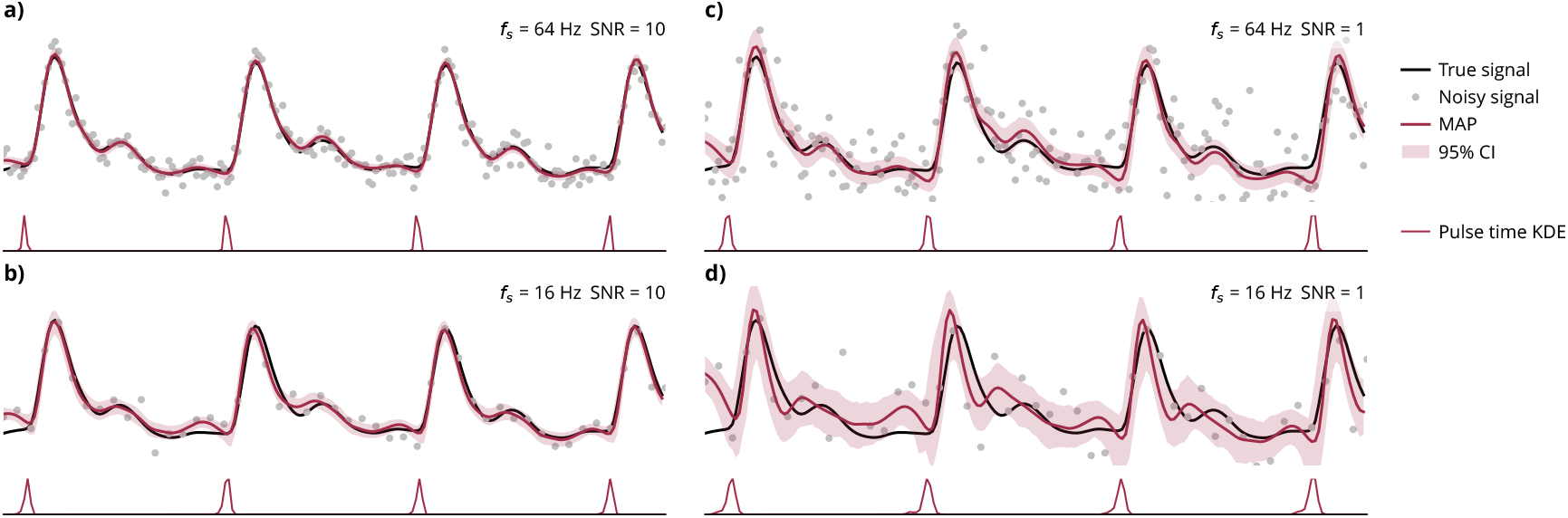
Uncertainty quantification. The true signal is derived from a 5-second photoplethysmogram waveform (only the first 3 seconds are plotted). Noisy signals are simulated with different sampling rates (*f*_*s*_) and signal-to-noise ratios (SNR). Shown in red are the maximum *a posteriori* (MAP) estimates and 99% credible intervals (CI) of the posterior were computed from 10,000 Monte Carlo Markov Chain (MCMC) samples. The marginal posterior distributions for the pulse times are shown as kernel density estimates (KDE) from the MCMC posterior samples. (**a**) High *f*_*s*_ = 64 Hz, high SNR = 10. (**b**) Low *f*_*s*_ = 16 Hz, high SNR = 10. (**c**) High *f*_*s*_ = 64 Hz, low SNR = 1. (**d**) Low *f*_*s*_ = 16 Hz, low SNR = 1.

## DISCUSSION

Simulation experiments demonstrate the advantages of joint signal filtering, timing detection, and shape extraction using Bayesian pulse deconvolution as compared to existing methods for vascular waveform analysis. Bayesian pulse deconvolution achieves better noise attenuation without introducing signal distortions, detects pulse timing with finer resolution, and extracts individual pulse shapes without truncating heart beats that overlap.

While simulations are required to evaluate algorithms against exact true values, they necessarily make simplifying assumptions that may not represent real vascular waveforms. It is important to consider how results in simulation generalize to human recordings. The simulated waveforms were designed to be as realistic as possible by sampling them from Bayesian prior densities derived from human recordings. Furthermore, the advantages of Bayesian pulse deconvolution persist when evaluated with human recordings. We conducted evaluations using the Aurora-BP dataset ^53^ because it is a large collection of raw vascular waveforms with precisely time-synchronized ECG, which is necessary for correctly evaluating timing accuracy. Precise pulse timing is important for accurately detecting arrhythmias and for accurately measuring high-frequency components of heart rate variability. ^8^ Future work should evaluate Bayesian pulse deconvolution on downstream applications that leverage the filtering and shape extraction features.

The Bayesian priors were derived from the same dataset of human recordings used to evaluate the pulse interval estimation accuracy. However, the Bayesian algorithm was not unfairly advantaged in the evaluation because the priors were fit using tonometry waveforms from a subset of individuals, while the evaluations were performed on photoplethysmogram waveforms from a non-overlapping subset of individuals.

The prior for the pulse shape coefficients **w** was fit to human recordings using pulse shapes extracted using the traditional segmentation and averaging method. To test whether pulse shape extraction error led to problematic distortions in the shape priors, we tested a prior bootstrapping method by which we used naive priors to perform Bayesian pulse deconvolution on the data and refit the shape priors on the maximum *a posteriori* pulse shapes. We found no significant changes in algorithm performance on our benchmarks, suggesting that the original data-driven priors were adequate.

Beyond improved performance, Bayesian pulse deconvolution offers other advantages. It quantifies uncertainty with a full joint posterior density over the inferred variables, which could be propagated to uncertainty in downstream derived metrics such as blood oxygen or blood pressure. Since it fits analytical signals to observations with arbitrary timing, Bayesian pulse deconvolution can be re-purposed to solve re-sampling, interpolation, imputation, and extrapolation tasks. These strengths could be significant for signals with sporadic windows of unusable data (e.g., wearable signals with motion artifacts), or adaptive sampling rates (e.g., power-optimized long-term continuous monitoring patches).

Another key advantage of the Bayesian framework is its extensibility to different modeling assumptions. For example, the assumption of constant average heart rate frequency could be relaxed to model sporadic arrhythmia events or respiratory heart rate variability. The model of pulse timing could be replaced by a temporal point process that better captures heart beat dynamics as measured by statistical goodness-of-fit. ^54,55^ The assumption of a static pulse shape could be relaxed to accommodate temporally varying pulse shapes. The pulse shape multivariate Gaussian prior could be replaced by a more flexible data-driven density estimate such as normalizing flows ^56^. The assumption of Gaussian noise could be modified to accommodate sensor-specific noise profiles or sporadic motion artifacts. Though this work considers only vascular waveforms measuring bulk flow in blood vessels, the method could be extended to cardiac waveforms (e.g., electro-cardiogram, phonocardiogram, ballistocardiogram, and seismocardiogram) which measure the excitation and contraction of the heart and have distinct morphologies. Additionally, the Bayesian priors could be personalized to individuals based on demographic factors or previous measurements.

While this work shows promise for Bayesian pulse deconvolution, the implementation presented here has some practical limitations. The challenge of multi-modality due to signal semi-periodicity scales combinatorially for signals with longer durations and greater pulse counts. This could be combated through buffered stochastic gradient Markov Chain Monte Carlo or variational inference approaches that scale inference in long time series. ^57^ Future work should allow pulse shapes to vary over time with per-subject or per-recording hierarchical modeling. Additionally, further optimization could improve the inference speed to enable real-time applications.

In light of these challenges, traditional vascular waveform analysis algorithms are in no way obsolete. Bayesian pulse deconvolution may be unwarranted for use cases that do not require extreme precision. For example, coarse estimates of heart beat intervals may be adequate to detect major arrhythmia events such as tachycardia, bradycardia, or atrial fibrillation. However, Bayesian pulse deconvolution may significantly impact applications requiring high timing precision (e.g., measuring autonomic modulation of heart rate), high noise reduction (e.g., remote photoplethysmography), or high shape extraction accuracy (e.g., non-invasive blood pressure estimation). Further work is needed to validate Bayesian pulse deconvolution for these and other downstream applications.

Nonetheless, the demonstrated benefits of Bayesian pulse deconvolution are significant and advance the state of the art for analyzing vascular waveforms. By jointly solving signal filtering, timing detection, and shape extraction in a unified algorithm, this method achieves superior performance compared with traditional sequential pipelines. This ability to extract more accurate insights from vascular waveforms could improve the performance of multiple clinical applications such as measuring autonomic function, non-invasively estimating blood pressure, measuring blood oxygenation, and designing sensitive, personalized biomarkers of cardiovascular health.

## METHODS

Bayesian pulse deconvolution aims to jointly solve three waveform analysis problems: filtering noise, detecting pulse timing, and extracting pulse shape. It achieves this by stipulating an analytical generative model that describes the supposed origin of the waveforms (**Fig. 6**). Unobservable variables are assigned Bayesian prior densities representing biologically and physically plausible values. Finally, inference is performed by updating a joint Bayesian posterior density over the unobserved variables conditioned on observed signal data. Analysis pipelines were implemented using Snakemake. ^58^

**Fig. 6.**
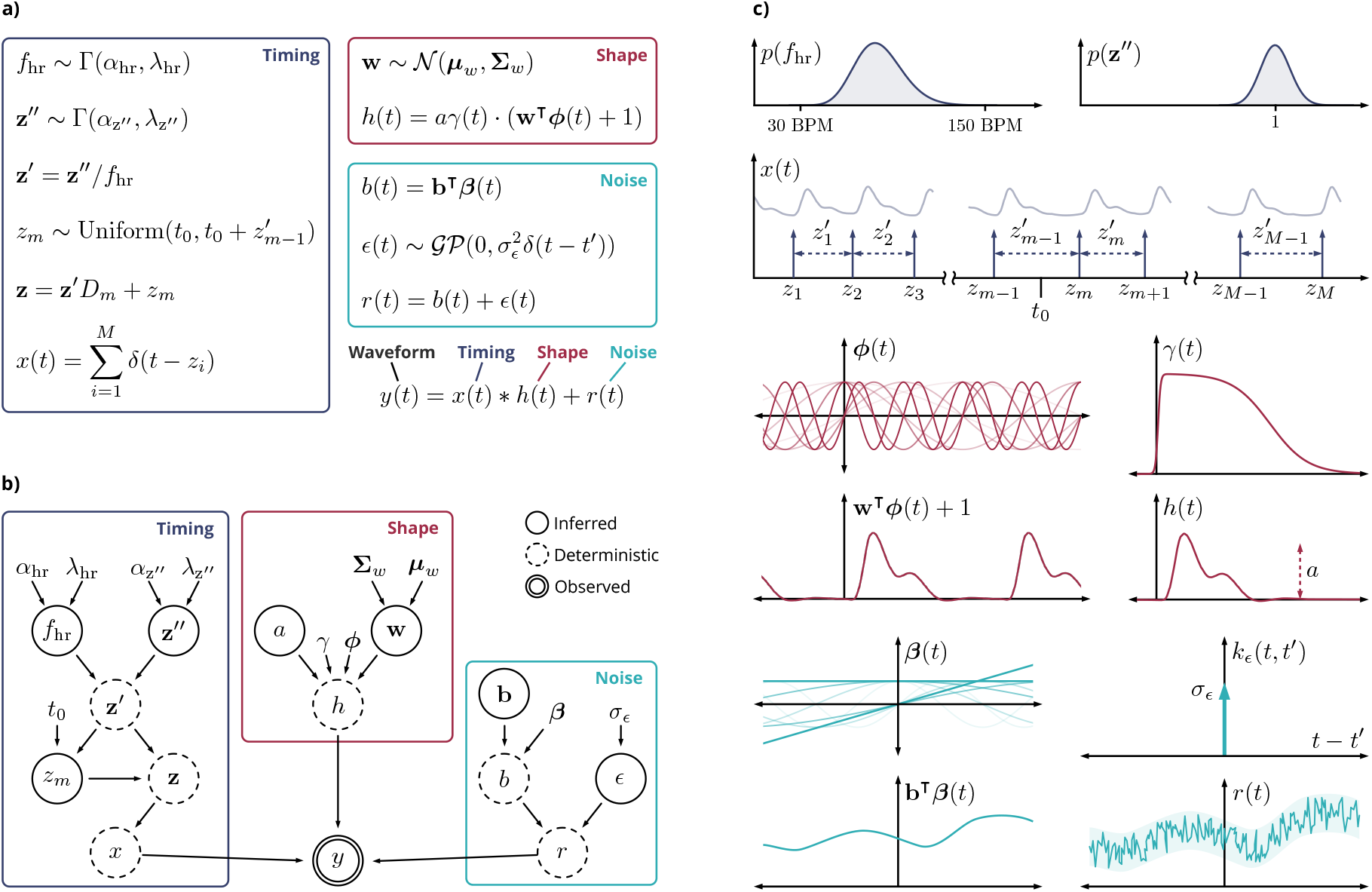
Analytical model of vascular waveforms. (**a**) Generative equations describing timing, shape, and noise components of observed signal *y*(*t*). (**b**) Bayesian graphical network representation. Arrows represent conditional dependence between variables. (**c**) Illustration of selected variables in the generative model.

### Analytical model of vascular waveforms

Vascular waveforms are described as a vector of samples **y** ∈ ℝ^*N*^ at sample times **t** ∈ ℝ^*N*^ (not necessarily uniform). These samples are modeled as observations of a continuous function of time *y*(*t*), where *y*_*i*_ = *y*(*t*_*i*_). The observable signal *y*(*t*) is modeled as a convolution of an impulse response *h*(*t*) with an impulse train *x*(*t*) plus a residual term *r*(*t*)

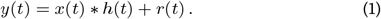

These terms correspond to the three goals of the algorithm. Filtering means isolating the true signal *x* ∗ *h* from the residual *r*, pulse timing is encoded by *x*, and pulse shape is encoded by *h*.

The residual consists of white noise *ϵ*(*t*) and a low-frequency baseline drift *b*(*t*). The white noise is modeled as a Gaussian process with zero mean *µ*_*ϵ*_(*t*) = 0 and i.i.d. covariance function 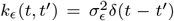, where *δ* is the Dirac delta function. The low-frequency baseline drift is modeled as the inner product of a weight vector **b** ∈ ℝ^*B*^ with a functional basis ***β***(*t*) = [1, *t*, sin(2*πf*_*b*_*t*), cos(2*πf*_*b*_*t*), sin(4*πf*_*b*_*t*), cos(4*πf*_*b*_*t*), …]^⊺^

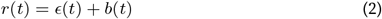

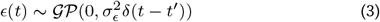

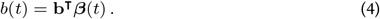

The impulse train *x*(*t*) is a sum of *M* Dirac delta functions *δ*(*t*) at the times of the pulse onsets **z** ∈ ℝ^*M*^

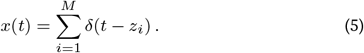

Initially, when the number of pulses is not known, *M* is initialized to be large enough that the pulse times **z** start before the first observed sample time *z*_1_ < *t*_1_ and end after the last observed sample time *z*_*M*_ > *t*_*N*_ . (Later, when the pulse count is known, *M* can optionally be reduced to trim unnecessary pulses outside the observed window, as described below.)

The impulse response *h*(*t*) represents the shape of a single pulse. It is constructed from a weighted combination of Fourier basis elements *ϕ*(*t*) = [sin(2*πf*_*h*_*t*), cos(2*πf*_*h*_*t*), sin(4*πf*_*h*_*t*), cos(4*πf*_*h*_*t*), …]^⊺^ with weights **w** ∈ ℝ^*K*^ . Since the Fourier components are periodic but the pulse shape is finite, we apply a temporally bounded window function *γ*(*t*). The pulse amplitude is determined by a constant scaling factor *a*

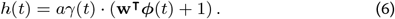

The window function is a product of two sigmoid functions, which supply left and right temporal bounds on the pulse shape

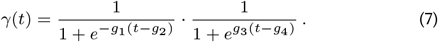

### Bayesian pulse timing priors

Priors on the vector **z** encode domain knowledge about pulse timing. This prior must satisfy two desired properties. Property 1: the prior should be time-shift invariant, that is, it should encode domain knowledge about the *relative* times between heart beats without making assumptions about the *absolute* times relative to the start of the observed signal. Property 2: the prior should avoid ambiguity or redundancy among values of *z* that differ only by cyclic permutations of the pulse times.

To address time-shift invariance (Property 1), we define the prior in terms of the relative inter-beat intervals. We define a pulse interval vector **z***′* ∈ ℝ^*M*−1^ encoding the times between sequential pulses

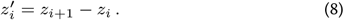

To address redundancy due to periodicity (Property 2), we arbitrarily assert that the middle pulse *z*_*m*_, where *m* = *M/*2, is the first pulse occurring after a time *t*_0_ in the center of the observed signal. That is, *z*_*m*−1_ < *t*_0_ < *z*_*m*_. Combined with Eq. 8, this assertion constrains the position of *z*_*m*_ to the range 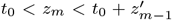. To preserve time-shift invariance, *z*_*m*_ has a uniform prior density on this range

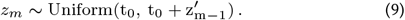

After sampling *z*_*m*_, all other pulse times *z*_1_, …, *z*_*m*−1_, *z*_*m*+1_, … *z*_*M*_ are fully determined by the pulse intervals **z***′*. The mapping from **z***′* to **z** is an affine transformation encoded by a matrix *D*_*m*_

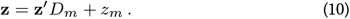

We assume that within a given observed signal there is a constant underlying average heart rate *f*_hr_ drawn from a gamma distribution

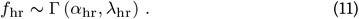

The gamma distribution shape and rate parameters *α*_hr_ and *λ*_hr_ were chosen so that 99.999% of the probability density falls between the physiologically realistic range of 30 BPM to 150 BPM.

Thus, the expected pulse interval is 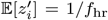. We use the heart rate *f*_hr_ to transform the pulse intervals **z***′* into a heart-rate-normalized form **z***′′* ∈ ℝ^*M*−1^ representing a percentage of the expected pulse interval

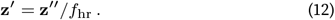

The normalized pulse intervals **z***′′* are assumed to be i.i.d. samples from a gamma distribution

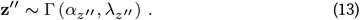

To fit the pulse interval parameters *α*_*z*_*′′* and *λ*_*z*_*′′*, we treat ECG R-R intervals as a pseudo-ground truth. These parameters are fit by maximum likelihood estimation to heart rate-normalized R-R intervals using 7,019 ECG recordings from 253 people in the Aurora-BP dataset. ^53^ To ensure 𝔼[**z***′′*] = 1, we applied the constraint that *λ*_*z*_*′′* = 1*/α*_*z*_*′′* .

### Bayesian pulse shape priors

The priors on the weights **w** encode domain knowledge about pulse shape. The weights are assumed to be multivariate Gaussian

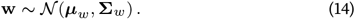

We assigned the shape distribution parameters ***µ***_*w*_ and **Σ**_*w*_ in a data-driven manner. The pulse shape prior was fit to 978 wrist-worn tonometry waveforms from 253 people in the Aurora-BP dataset. ^53^ Tonometry signals were chosen because they contain a higher frequency bandwidth than photo-plethysmogram signals. The average segmented pulse shapes from the waveforms were used to fit the window function parameters (*g*_1_, *g*_2_, *g*_3_, and *g*_4_) and 978 weight vectors **w** using maximum likelihood estimation with Eq. 6. The parameters ***µ***_*w*_ and **Σ**_*w*_ were then fit to the data-derived weight vectors **w** using maximum likelihood estimation.

Since pulse segmentation is not a ground truth method for extracting pulse shapes from vascular waveforms, we took measures to mitigate bias in the data-driven prior. To mitigate filtering distortions, the waveforms were not filtered; instead, we selected only the waveforms with the highest signal quality score assigned by the dataset creators. ^53^ To mitigate error introduced by inexact pulse timing estimates, we used time-synchronized ECG R-R intervals as pseudo-ground truth pulse times. Finally, to mitigate pulse truncation due to overlapping pulses, we selected only recordings with long beat-to-beat intervals (heart rates less than 55 BPM).

### Bayesian inference

The Bayesian inference algorithm was implemented using the NumPyro probabilistic programming package, ^59,60^ which is built on the JAX hardware acceleration library. ^61^ We used the optax library for learning rate scheduling. ^62^ Maximum *a posteriori* (MAP) solutions were estimated using stochastic variational inference with a Dirac delta approximate posterior form. We use the reparameterization trick to improve efficiency of sampling and gradient-based optimization. ^63^ The scaling factor *a*, offset weights **b**, and white Gaussian noise variance *σ*_*ϵ*_ are all unconstrained parameters allowed to vary freely during inference.

The posterior distribution has multiple modes corresponding to solutions with missing or extra pulses. Since the total number of pulses *M* is unknown *a priori*, we first choose *M* so that the estimated pulse times **z** extend before and after the observed time points. To efficiently converge on the mode with highest posterior, we initialize the MAP estimation with multiple states and keep only the mode with the highest posterior density. Initial states are sampled from the prior, with the heart rate *f*_hr_ heuristically biased by the signal’s power spectrum. After locating the mode, the pulse count *M* is decreased to trim pulses outside the observed time range.

### Uncertainty quantification

For uncertainty quantification, we first used Bayesian pulse deconvolution to fit an analytical waveform to a real photoplethysmogram waveform. For illustrative purposes, we used this analytical solution to simulate four 3-second noisy waveforms with different sampling rates and signal-to-noise ratios. To quantify the model uncertainty we drew 10,000 posterior samples using Markov Chain Monte Carlo No-U-Turn sampling. ^64^

### Baseline algorithms

We evaluated Bayesian pulse deconvolution relative to representative vascular analysis algorithms reported in the literature and available in widely used Python packages (**Table 1**).

### Simulation evaluations

Simulated vascular waveforms were generated by sampling from the empirical Bayesian priors, with unit scaling factor *a* = 1, zero baseline offset **b** = 0. The waveforms were sampled in 11-second windows with a uniform 64 Hz sampling rate and a noise level *σ*_*ϵ*_ tuned to –20 dB of the true signal power. These sampling parameters were intended to represent realistic vascular waveforms. To confirm the generalizability across sampling parameters, we performed parameter sweeps across sample durations, sampling rates, and noise levels **(Sup. Fig. 1)**.

To evaluate filter accuracy, we compared the pre-filter error (observed signal minus true signal) and the post-filter error (filtered signal minus true signal); we then computed decibel power attenuation between the pre- and post-filter error. To mitigate error due to edge effects from traditional infinite impulse response filters, signal power was weighted by a Hann function spanning the signal duration. To isolate only error due to changes in the relative waveform contours, we scaled all signals to unit mean and variance. To evaluate pulse timing accuracy, we first matched estimated pulse times with their corresponding true pulse times, discarding false positives and false negatives. Within each waveform, we computed the root-mean-square error between estimated and true pulse intervals.

To evaluate pulse shape extraction, we developed a metric that quantifies extracted pulse shape error as a function of the total area of the true pulse shape. To compute this metric, we first aligned the algorithms with a time shift that maximized the Pearson correlation between true and predicted pulse shapes. We used zero-padding to standardize pulse shapes of different lengths. We then scaled both signals to have unit area (integrating to one). The error metric was defined as the total unsigned area of the difference between the true and predicted pulse shapes.

### Evaluations on human recordings

We evaluated Bayesian pulse deconvolution using the Aurora-BP dataset which includes raw photoplethysmogram waveforms and time-synchronized ECG signals. ^53^ Since irregular motion artifacts are out of scope for this work, the evaluation was performed only on participants with high heuristic optical signal quality indices, as assigned by the dataset authors. We measured gold standard pulse intervals using ECG R-R intervals and quantified agreement with predicted pulse times using root-mean-square error.

### Statistical analysis

All algorithms were statistically compared pairwise using Wilcoxon signed-rank test with a two-sided alternative hypothesis. To account for multiple hypothesis testing, Bonferroni corrections were applied to all p-values within each figure. Not all statistically significant differences are annotated.

## DATA AVAILABILITY

The dataset of simulated waveforms generated for this work are available from the authors upon request. The Aurora-BP dataset is available from its authors upon request, with information available at https://github.com/microsoft/aurorabp-sample-data.

## CODE AVAILABILITY

The code used to implement Bayesian pulse deconvolution, generate simulated vascular waveforms, run analyses, and generate figures in this work is available at https://doi.org/10.5281/zenodo.18586098.

## ACKNOWLEDGMENTS

We thank Michael Li, Ron Polonsky, Eli Waldman, Alison Marsden, and John Duchi for their insights and feedback.

This work was supported in part by the Wu Tsai Human Performance Alliance at Stanford University, the Wu Tsai Neurosciences Institute at Stanford University, the Stanford Bio-X Interdisciplinary Biosciences Institute, and the Stanford Institute for Human-Centered Artificial Intelligence (HAI). EBF is a Chan Zuckerberg Biohub – San Francisco Investigator.

This work is supported in part by NSF awards BCS-1932619 and 2205084, ONR grant N00014-22-1-2110, and NIH awards F31HL176112, U01NS131914, and 1U01DK140939. The content is solely the responsibility of the authors and does not necessarily represent the official views of the National Institutes of Health and a Predoctoral Individual National Research Service Award from the National Institutes of Health.

## AUTHOR CONTRIBUTIONS

P.S.R., J.A.L., T.P.C., and E.B.F. conceptualized the algorithm.

P.S.R., T.D., and L.O. implemented and evaluated the algorithm.

P.S.R., T.D., and L.O. drafted the manuscript and figures.

J.A.L., E.B.F., and T.P.C. read and revised the manuscript.

