## Supplementary material for "Vascular waveform analysis using Bayesian pulse deconvolution": Sup. Fig. 1

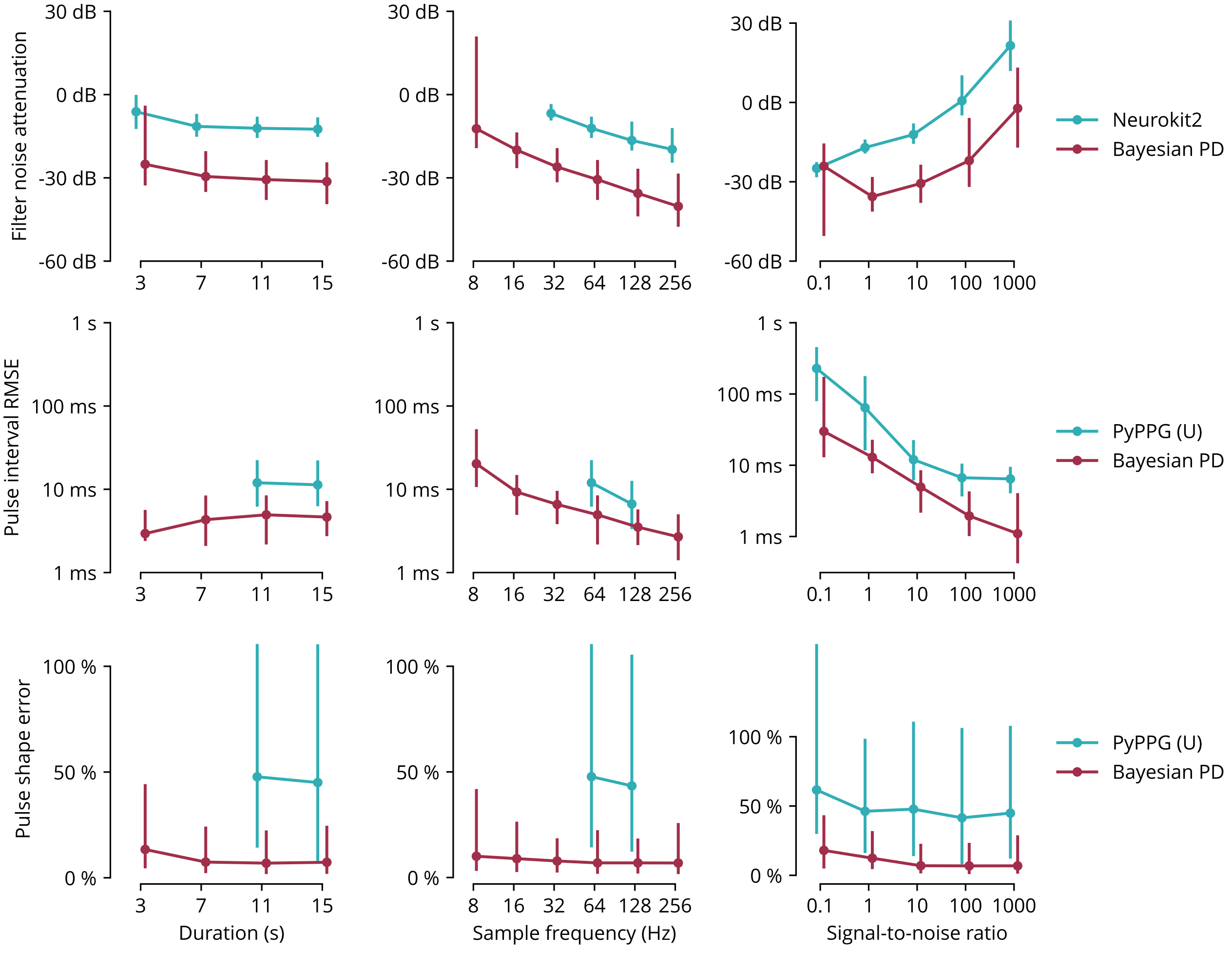


**Supplementary Fig. 1 | Algorithm performance across sampling parameters.** We compared Bayesian pulse deconvolution (PD) and the best-performing existing method across different signal durations, sampling frequencies, and signal-to-noise ratios. In all conditions, Bayesian pulse deconvolution achieves lower noise attenuation, lower root-mean-square error (RMSE) of pulse interval timing, and lower percent area error in pulse shape. Error bars represent 90th percentiles out of 100 simulated waveforms for each condition. Baseline algorithms are not plotted for conditions that are incompatible with their Python package implementations.
